# A structural antibody benchmark for leakage-aware deep-learning evaluation

**DOI:** 10.64898/2026.09.06.749672

**Authors:** Tomer Cohen, Haimasree Bhattacharya, Michal Ozery-Flato, Dina Schneidman

## Abstract

Deep-learning methods for antibody structure prediction, antibody-antigen interaction modelling and design are advancing rapidly. However, comparisons across studies remain difficult because training and test sets are often constructed independently, and a temporal cutoff alone does not prevent train-test leakage. We present SABLE (Structural Antibody Benchmark for deep-Learning Evaluation), a versioned structural antibody resource that couples a fixed training collection with a leakage-controlled held-out test set for reproducible machine-learning development and evaluation. SABLE combines 16,511 experimental training entries with 327 manually reviewed test entries and 3,274 high-confidence, patent-derived AlphaFold3 models spanning 476 antigens. Candidate test structures were selected after the AlphaFold3 temporal cutoff and filtered against the training set using antibody and antigen sequence similarity filters. Each test entry records its nearest training set neighbour, allowing users to quantify remaining relatedness and stratify performance by similarity. Versioned releases provide metadata, processed structures, CDR annotations, redundancy labels and model-confidence fields. A Python/PyTorch API and standardised benchmark metrics provide reproducible database access and evaluation code without requiring additional antibody-structure processing.

## Background & Summary

Antibodies are central mediators of adaptive immunity and an important class of therapeutic biomolecules. Their antigen-binding site is formed by two protein chains, heavy and light. Each chain contains a relatively conserved framework region and three highly variable and diverse loops, known as complementarity-determining regions (CDRs). Together, the six CDRs account for much of the sequence and structural diversity among antibodies, with the third CDR of the heavy chain (H3) being the most variable in sequence, length and conformation. Due to this architecture, antibodies are challenging targets for computational modelling^1–8^ and design^9–18^. Antibody-antigen complex prediction is further complicated because antibodies and their antigens share no evolutionary history, so paired multiple-sequence alignments carry little information about how they bind^1,2,19–21^. Moreover, antibody binding sites (epitopes), while not uniformly distributed on antigens, can occur across diverse antigen surfaces and geometries^22^. Consequently, despite major advances in general protein structure prediction, accurate CDR modelling, antibody-antigen docking, and ranking of candidate complexes remain challenging^1,2,23,24^.

Reliable progress in this setting depends not only on method design but also on how training and evaluation data are constructed. Established machine-learning fields address this with fixed benchmarks that define common training and test splits, so that reported gains can be attributed to the method rather than to differences in data. ImageNet, for example, has served this role in computer vision, and GLUE in natural language processing^25,26^. Antibody modelling has instead generally relied on study-specific snapshots, data filtering rules, and test sets. Direct comparison between the resulting methods is difficult, because studies differ in the antibody regions used to measure similarity; the sequence-identity thresholds applied; whether antigen similarity is considered; and the structures and dates used to define training and test sets. Even when the same data aíe used, evaluation procedures can differ. For example, reported CDR RMSDs depend on the CDR definition and the residues used for framework superposition, while antibody-antigen DockQ^27,28^ calculations can depend on chain selection and complex representation. These differences make it difficult to determine whether an apparent performance improvement reflects a better model, a less challenging test set, differences in data overlap, or differences in the implementation of evaluation metrics.

Several complementary resources provide the experimentally determined structures of antibodies and antibody-antigen complexes, from which antibody-specific machine-learning datasets can be constructed. The Protein Data Bank (PDB) is the global archive of experimentally determined macromolecular structures and provides the underlying structural records used by most antibody structure resources^29^. SAbDab adds antibody-specific curation to PDB structures, including heavy- and light-chain assignments, antigen annotations, antibody numbering and downloadable processed structures^30,31^. SAbDab2 extends this resource with enhanced antibody-level identifiers, regular updates and versioned train/test partitions intended for machine-learning use^32^. SAAINT-DB independently derives antibody and antibody-antigen annotations from PDB structures, with an emphasis on parsing coordinate files, chain pairing, interaction identification, and curated binding-affinity information^33^. These structural databases are valuable sources of antibody data, but using their records in reproducible machine-learning benchmarks still requires explicit choices about temporal training and test set cut-offs, redundancy thresholds, sequence similarity metrics, treatment of missing coordinates, variable-domain boundaries, CDR definitions, and antigen representation.

Other resources cover complementary antibody data that lacks experimental structures. Observed Antibody Space (OAS) collects and annotates large-scale antibody repertoire sequencing data and contains more than a billion antibody sequences. Some records contain paired heavy and light chain sequences, but neither their structures nor their antigen targets are known^34^. PLAbDab integrates paired antibody sequences extracted from patents and the scientific literature, links them to source documents and functional annotations, and provides predicted antibody structures for structural searching^35^. The NaturalAntibody Patents Database similarly mines large collections of patented antibody sequences and connects antibody records with antigen target and UniProt annotations^36^. Resources focused on interaction measurements address yet another need: PPB-Affinity aggregates experimentally determined protein-protein complex structures with binding-affinity annotations^37^. FLAb provides a complementary benchmark of experimentally measured antibody fitness landscapes across properties including expression, thermostability, immunogenicity, aggregation, polyreactivity and binding affinity^38^. Antibody sequence resources therefore cover a sequence space far larger than that represented by experimentally determined structures, but they lack the structural information that can facilitate machine learning.

The difficulty of predicting antibody-antigen structures has motivated several dedicated benchmarks, alongside general biomolecular benchmarks that include antibody-antigen complexes. Fromm, Ludaic and Elofsson constructed a benchmark of 110 antibody-antigen complexes released post-September-2021 from SAbDab, filtered against pre-cutoff structures using an 80% sequence-identity threshold applied separately to heavy, light and antigen chains, and subsequently evaluated several state-of-the-art structure-prediction methods^39^. Hitawala and Gray assembled an antibody and nanobody (a single-chain antibody that lacks a light chain) benchmark to examine AlphaFold3 and related approaches using antibody structural accuracy metrics, including H3 RMSD, together with antibody-antigen docking quality^40^. FoldBench provides a broader all-atom benchmark of 1,522 low-homology targets across nine biomolecular prediction tasks, including 172 antibody-antigen interfaces^41^. Yu et al. compared antibody structure prediction, antibody-antigen docking and interface analysis methods under a common controlled evaluation design^42^. Continuously updated evaluation initiatives, such as CoFold Arena, provide public leaderboards for biomolecular co-folding methods^43^. These benchmarks are valuable for assessing existing predictors, but they provide only test sets for evaluation, without a corresponding fixed training set, and a release-date cutoff alone does not ensure that test entries are dissimilar to what a method was trained on. A complete antibody-specific development resource that integrates fixed training and test sets, data leakage controls, per-target relatedness metadata, and shared evaluation software could further enhance model training and evaluation.

SABLE complements existing antibody resources by integrating curated structural data with an explicitly defined framework for machine-learning training and evaluation. To construct a leakage-controlled test set, the experimental dataset applies the commonly used AlphaFold3 training-data cutoff of 30 September 2021, followed by antibody- and antigen-level sequence-similarity filters, structural-quality criteria and manual review. The resource further includes high-confidence AlphaFold3-modelled antibody-antigen complexes derived from patented antibody sequences, extending coverage to antibody-antigen pairs without experimentally determined structures. Importantly, every test entry is annotated with its closest experimental and modelled training neighbours according to H3 distance, variable-domain sequence identity and, where applicable, antigen sequence similarity. Versioned data releases are accompanied by a GitHub repository, Python/PyTorch API, and web interface providing access to sequences, annotations and structures, together with implementations of antibody-specific sequence similarity and structural-accuracy measures, including H3-distance, sequence-identity, framework-aligned CDR RMSD and DockQ calculations. Collectively, these resources provide a reproducible basis for developing and evaluating antibody-focused machine-learning methods.

## Methods

### Data sources

Experimental structure records were derived from the SAbDab snapshot downloaded on 29 April 2026. SAbDab annotates antibody structures deposited in the PDB with heavy-chain, light-chain and antigen-chain assignments and associated metadata^29–31^. The source table included PDB identifiers, release dates, experimental methods, resolution and crystallographic R values where available, antibody format, species and subclass annotations, antigen names, types and species, and affinity-related fields where present. A total of 21,023 source SAbDab rows yielded 20,279 processable antibody entries.

The patent-derived sequence data originated from the NaturalAntibody Patents Database snapshot of 30 July 2025^36^.

The release described here is SABLE v0.2.0, deposited in Zenodo under DOI 10.5281/zenodo.21759053.

### Processing of experimental antibody structures

#### Entry definition and chain selection

A single PDB file can yield more than one SAbDab annotation row. Each row was processed independently, and the stable experimental identifier was formed from the lower-case PDB identifier and the row index within that PDB (for example, “7myt_0”). The SAbDab row and its chain assignments defined the molecular unit used by SABLE. The first coordinate model in the referenced PDB file was read with Bio.PDB^44^. SAbDab metadata was used to select the antigen and antibody heavy and light chains. For single-chain Fv records in which the heavy and light variable domains share a physical chain, the two domain intervals were separated using their annotated sequences and numbering.

#### Variable-domain and CDR annotation

Standard and modified amino-acid residues were converted to one-letter sequences, and AbNumber was used with the IMGT CDR definition to identify the variable domain and its framework (FR1-FR4) and CDR1-CDR3 regions^45,46^. The release stores complete heavy- and light-chain variable-domain sequences, CDR sequences, and zero-based half-open (start, end) intervals locating each region in its parent sequence. Coordinate renumbering was not applied; processed PDB files retain source chain identifiers, residue numbers, and insertion codes.

#### Missing, modified and unknown residues

PDB REMARK 465 records were used to identify missing residues that are not present in the structure. Missing positions are retained in antibody and antigen sequences as lower-case one-letter symbols and have false coordinate masks. The fields “has_missing_in_cdrs”, “max_missing_gap” and “sum_missing” summarise coordinate incompleteness of the antibody chains. Modified amino acids are mapped to their canonical one-letter parents in sequence-related metadata fields, and unknown amino acids are represented as X.

#### Antigen extraction and sequence representation

SAbDab-assigned antigen chains were retained in the processed structure after waters, common crystallisation additives, and simple ions were removed. Protein and peptide chains are represented by one-letter amino-acid sequences; nucleic-acid chains by one-letter nucleotide sequences; and haptens and carbohydrates by ordered PDB Chemical Component Dictionary (CCD)^47^ codes separated by hyphens. Antigens consisting of multiple chains have their sequences concatenated using “|” as a delimiter.

### Processing of synthetic patent-derived antibodies

The synthetic dataset originated from an initial 217,463 NaturalAntibody patent-derived records. Because NaturalAntibody did not contain heavy-light pairing for most entries, pairing was recovered by comparison with the PLAbDab snapshot of 24 November 2025^35^. Nanobody records were retained directly. For paired antibodies, a NaturalAntibody heavy chain was assigned a PLAbDab light chain when the internal heavy-chain sequence, after removing three residues from each terminus, was contained in the comparison sequence in either direction. This pairing procedure produced 184,160 antibody-antigen pairs with sequence records representing 26,415 antibodies and 5,090 antigens. Antigen UniProt identifiers were taken from the NaturalAntibody metadata, and the corresponding sequences were retrieved from UniProt^48,49^. Antigen sequences longer than 1,000 amino acids were excluded, leaving 165,230 candidate complexes. AlphaFold3 was used to model all candidate complexes. Multiple sequence alignment generation used a reduced antibody-specific sequence database and antigen-specific databases were generated with Jackhmmer^50,51^. One random seed was used to generate five model samples per candidate complex. Confidence selection used an antigen-focused interface predicted TM (ipTM) score, so that the heavy-light interface did not dominate the complex-level confidence measure. For each candidate complex, the highest-ranked of the five samples was retained if its antigen-focused ipTM was ≥ 0.8, yielding 5,843 high-confidence structures.

### Train-test split protocol

#### Initial temporal and quality split

Experimental structures released on or before 30 September 2021 formed the initial training pool, whereas later structures formed the candidate held-out test pool. The date was selected to align with the commonly used structural-data cutoff reported for AlphaFold3^52^ as well as OpenDDE^53^, OpenFold^54^, Boltz-1^55^ and other tools. Post-cutoff entries were grouped together if they had the same sorted antigen-type combination and the exact six-CDR sequence; the highest-quality representative of each group was selected using the lexicographic tuple of numerical structure resolution, presence of missing CDR residues, and total missing-residue count. Candidate test entries were required to have a reported resolution of 3.0 Å or better and no missing residues in annotated CDRs. 459 light-chain-only entries were excluded before split construction.

#### Antibody sequence similarity filtering

Antibody similarity was calculated with a global Biopython pairwise aligner configured with match = 1, mismatch = 0, and gap = 0^44^. Under these scores, the alignment score equals the longest-common-subsequence (LCS) length.

Two measures were used to filter the candidate test pool against the training set: H3 distance and variable-domain identity.

H3 distance between two loops (*a*, *b*) was defined as an LCS-derived dissimilarity:

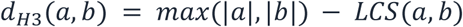

The H3 sequence of every retained test entry had a distance of at least 3 from the H3 sequence of every experimental training entry. Variable-domain identity was defined as 100 × LCS length divided by the shorter sequence length. When both entries contained light chains, SABLE used the larger of the heavy-chain-only identity and the identity of the concatenated heavy-and light-chain sequences. Retained test set entries had variable-domain identity no greater than 90% to every experimental training set entry. An H3 distance of at least three excludes exact and near-exact H3 matches, while the 90% variable-domain similarity ceiling additionally removes antibodies that remain highly similar outside H3.

#### Antigen sequence similarity filtering

Protein- and peptide-containing antigens were screened with BLASTP^56^. Initial-training sequences were written to a protein BLAST database, and candidate-test antigens were queried using an E-value ceiling of 10, up to 100,000 target sequences, and tabular output containing full query and subject lengths. Query, subject, and shorter-sequence coverage were calculated as 100 × alignment length divided by the corresponding full-sequence length or the shorter full-sequence length. A candidate was excluded if any hit had percent identity of at least 30%, shorter-sequence coverage of at least 70%, and E-value of at most 1 × 10^-5^, or if any hit had percent identity of at least 70% irrespective of coverage and E-value. The first criterion was used to identify substantial sequence-level relatedness, whereas the unconditional 70% identity rule captures highly similar local regions even when full-length coverage is limited. Entries without a structure-derived protein or peptide antigen sequence were not subjected to the automated BLAST filter.

#### Enriching the training set with failed test candidates

Candidate test records satisfying the temporal, structural-quality, antibody-similarity and antigen-similarity criteria formed a provisional test set. A quality-failing group representative and all members of a CDR/antigen-type group whose representative failed a similarity rule entered an add-back to the training set pool. These entries were compared with the provisional test set using the same antibody and antigen criteria. Only entries without detected overlap with the provisional test set were added to the training set. Entries that remained too similar were excluded from both components, and non-representative members of groups represented in the test set were not added to training. The automated workflow produced 338 candidate test entries.

#### Manual inspection

To ensure the quality of the held-out test set, all candidate test entries were manually inspected for antibody and antigen chain assignments, antigen type, structural representation, antigen stoichiometry, and metadata consistency. Manual inspection identified 24 entries requiring correction or exclusion (Table S1). Eleven entries were removed because the processed record did not provide a sufficiently unambiguous representation for benchmark evaluation, including non-canonical or fused antibody architectures such as diabodies, megabodies, and light-chain-nanobody complexes; incorrect or ambiguous antigen annotations; and duplicate or less appropriate representations of another retained complex. The remaining 13 flagged entries were retained after manual correction, primarily of antigen-chain assignments and multimeric antigen representation (Table S1). This inspection yielded 327 final test entries.

#### Synthetic dataset filtering

Two overlap filters were applied to the patent-derived modelled data after the final experimental training and test sets had been established. First, a modelled entry was removed when its heavy- and light-chain variable-domain sequences each exactly matched, or were contained within, an experimental-training entry, or when all six CDR sequences matched an experimental-training entry. Second, the remaining modelled entries were compared with the final held-out test set and removed if they failed the antibody or antigen leakage-control rules defined above. The final modelled component contains 3,274 complexes covering 476 antigens.

#### Test-set nearest-neighbour metadata

For every test entry, the release records the nearest experimental-training and patent-derived modelled neighbours by minimum H3 distance and maximum variable-domain identity. For protein or peptide antigens, the stored BLAST neighbour is selected by descending bit score, then descending percent identity and shorter-sequence coverage, and finally ascending E-value. Accordingly, the stored percent identity corresponds to the best-ranked BLAST hit and is not necessarily the maximum percent identity among all BLAST alignments.

### CDR redundancy groups

To make redundancy within the experimental training collection explicit and support flexible dataset construction, entries were assigned to groups based on identical CDR sequences. After the final experimental training collection was established, entries were grouped by the exact ordered six-CDR tuples (H1, H2, H3, L1, L2, and L3). Heavy-chain-only records have empty light-chain CDR fields. Each cluster receives the identifier of its selected representative in “redundancy_cluster”, and that record is marked “non_redundant=True”. Representatives were selected deterministically using the same quality tuple as above: lower numerical resolution, absence of missing CDR residues, then fewer missing residues. This allows users to train models on either the complete redundancy-preserving experimental collection or one with a selected representative per exact-CDR group.

### GitHub and web server interfaces

The accompanying GitHub repository provides code for retrieving and using the dataset. The Python package downloads the fixed Zenodo record, verifies checksums, performs path-safe archive extraction, reconstructs a consistent local release tree and validates required fields and structure paths. “SableDataset” and “TorchSableDataset” expose composable filters for split, experimental/modelled origin, redundancy, and antigen availability. Metadata-indexed coordinate helpers retain unresolved sequence positions as zero coordinates with false masks, preventing downstream coordinate tensors from shifting after missing residues.

Each dataset entry is represented by an “Antibody” object that combines its metadata with access to the corresponding processed structure. The object provides direct access to heavy- and light-chain sequences, framework and CDR sequences and indices, antigen annotations and sequences, model-confidence and nearest-neighbour fields, and the underlying antibody and antigen residues. Structure-aware methods provide metadata-indexed Cα or selected-atom coordinates and masks for unresolved positions, as well as antigen coordinates, antibody-antigen interface residues and evaluation metrics such as framework-aligned CDR RMSD and DockQ^27^. Thus, common antibody-specific operations can be performed without requiring users to implement PDB parsing, CDR indexing, missing-residue handling or antigen-chain selection themselves.

The web server reads the same release and provides separate views for experimental training entries, patent-derived modelled entries and held-out test entries. Users can search metadata, inspect sequences and CDR annotations, visualise processed structures and follow links to the Zenodo release and software repository. Figure 1 summarises the complete data-construction and access workflow.

**Figure 1.**
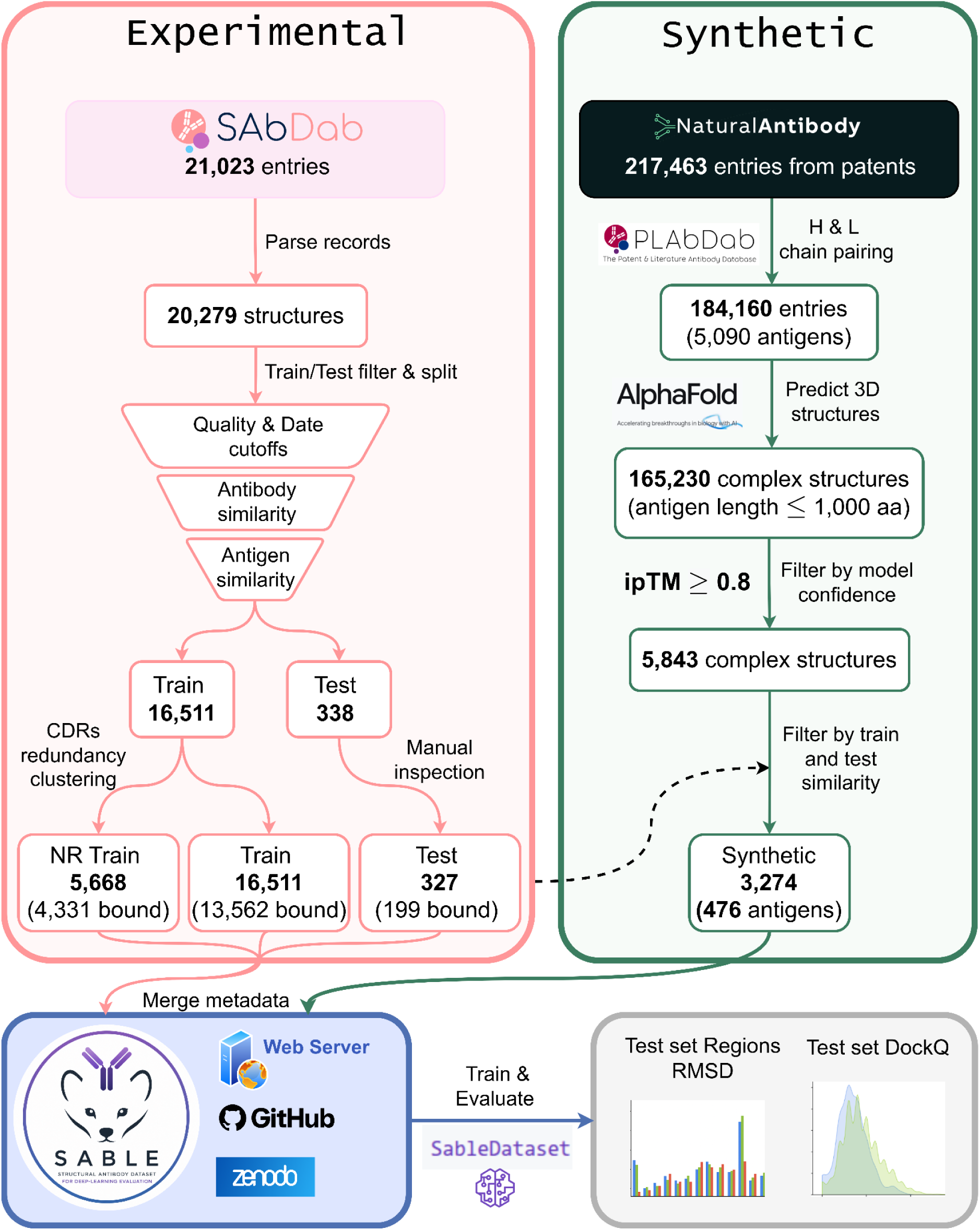
Overview of dataset construction and access. Experimental antibody annotations from SAbDab were parsed into processed variable-domain structures, separated by release date, reduced to quality-ranked representatives for candidate testing, filtered for antibody and antigen similarity, and manually reviewed to obtain 327 held-out entries. The final experimental training set contains 16,511 entries, including 5,668 exact-CDR representatives. Patent-derived NaturalAntibody records were paired with light chains using PLAbDab, filtered to antigens no longer than 1,000 residues, modelled with AlphaFold3, retained at antigen-focused interface confidence “antigen_iptm” ≥ 0.8, and filtered against the experimental training and test sets to obtain 3,274 complexes covering 476 antigen targets. Counts correspond to SABLE v0.2.0. The three components share one schema and are distributed through Zenodo, the web server, GitHub and the Python/PyTorch API, which also provides benchmark-evaluation utilities including H3 RMSD and DockQ.

## Data Records

SABLE is deposited in Zenodo under DOI 10.5281/zenodo.21759053^57^. Zenodo contains version-prefixed files so that individual objects can be downloaded and verified independently. The package reconstructs these flat files into a local release tree. Table 1 summarises the deposited objects.

**Table 1.** Files in the SABLE v0.2.0 Zenodo release.

| File or pattern | Format | Contents |
| --- | --- | --- |
| sable_v0.2.0_metadata.csv and .parquet | CSV; Apache Parquet | Equivalent per-entry metadata in human-readable and typed/high-throughput forms. |
| sable_v0.2.0_metadata_schema.json | JSON | Machine-readable field names, nullable types and constraints. |
| sable_v0.2.0_column_descriptions.md | Markdown | Human-readable definitions for all metadata columns. |
| sable_v0.2.0_splits.zip | ZIP; text; JSON | Predefined train/test identifiers, experimental/modelled subsets and redundancy mappings. |
| sable_v0.2.0_sequences.zip | ZIP; FASTA | Heavy-chain, light-chain and paired variable-domain sequences. |
| sable_v0.2.0_structures_pdb_*.tar.gz | Gzip-compressed tar; PDB | Sharded processed structure files, one `structure_path` per entry. |
| sable_v0.2.0_manifest.json | JSON | Release inventory, file roles and extraction destinations. |
| sable_v0.2.0_checksums.sha256 | Text | SHA-256 hashes for deposited files. |
| sable_v0.2.0_VALIDATION_REPORT.md | Markdown | Release-construction and consistency validation summary. |
| sable_v0.2.0_LICENSE-DATA.txt and README.md | Text; Markdown | Data-use terms and release documentation. |

Each metadata row corresponds to one processed antibody entry. “entry_id” is the stable primary key, and “structure_path” locates its PDB file. “split” takes the value “train” or “test”. The boolean “synthetic” distinguishes patent-derived modelled structures from experimental structures, while “non_redundant” marks the selected representative of an experimental “redundancy_cluster”. These fields are independent: an experimental training row may be redundant or non-redundant, whereas every modelled row remains explicitly identified even though it belongs to the training split.

The metadata contains 97 released fields. Table 2 groups representative columns. The complete column dictionary and JSON schema define field types, missing-value rules and aligned multi-chain fields. Held-out test rows store nearest experimental-training and modelled neighbours in the “closest_*_h_cdr3_distance”, “closest_*_seq_identity” and “closest_*_blast_*” fields.

**Table 2.**
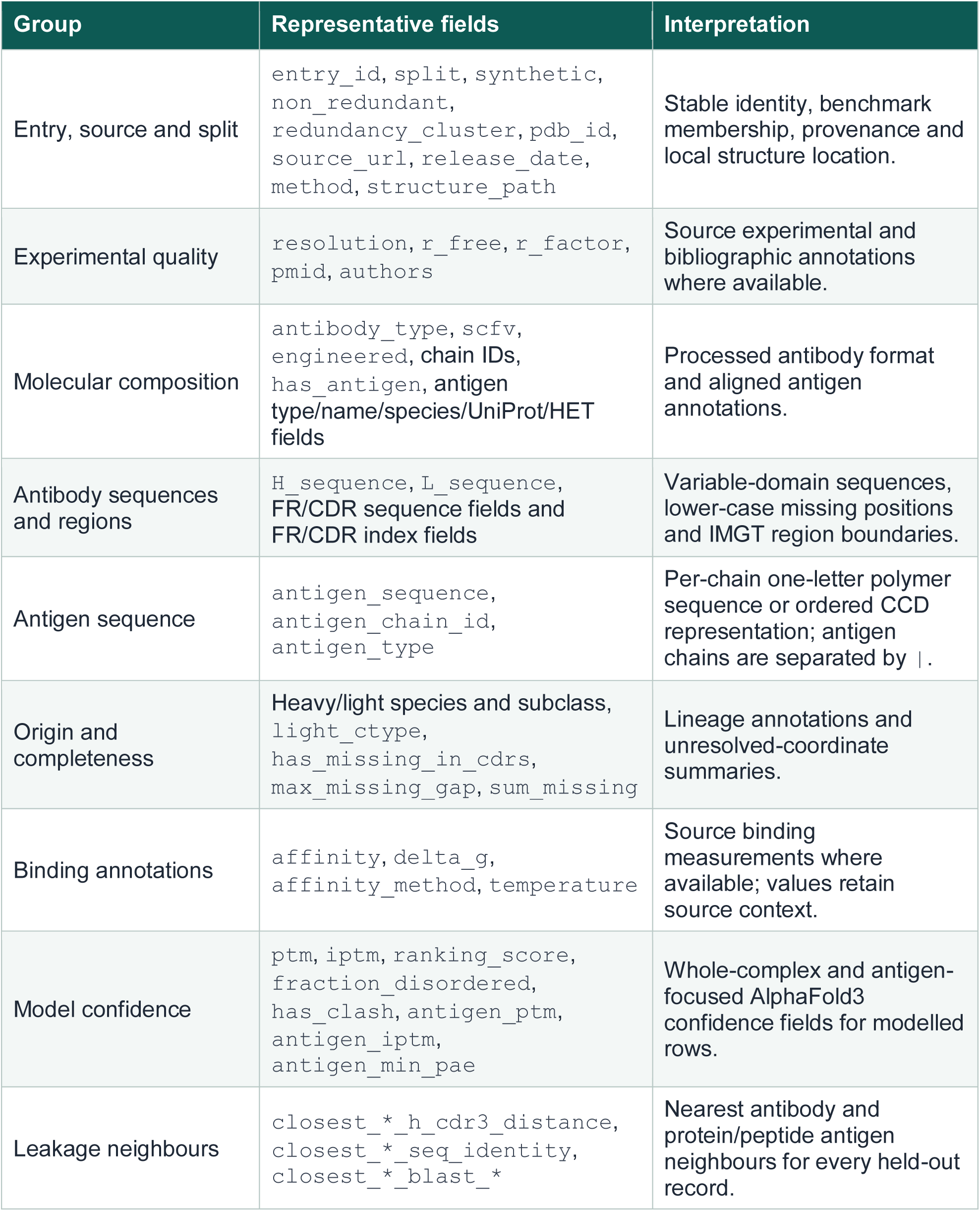
Metadata groups and representative fields.

After extraction with the SABLE software, the release has the following directory structure:

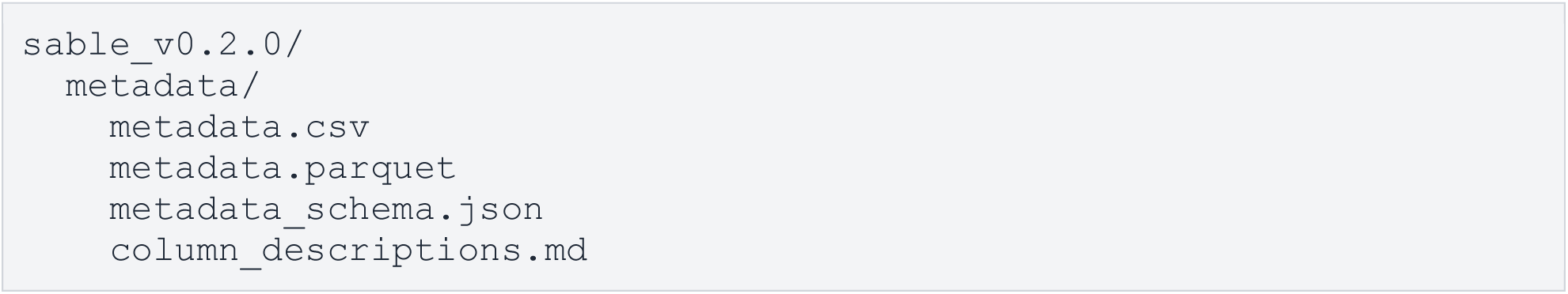
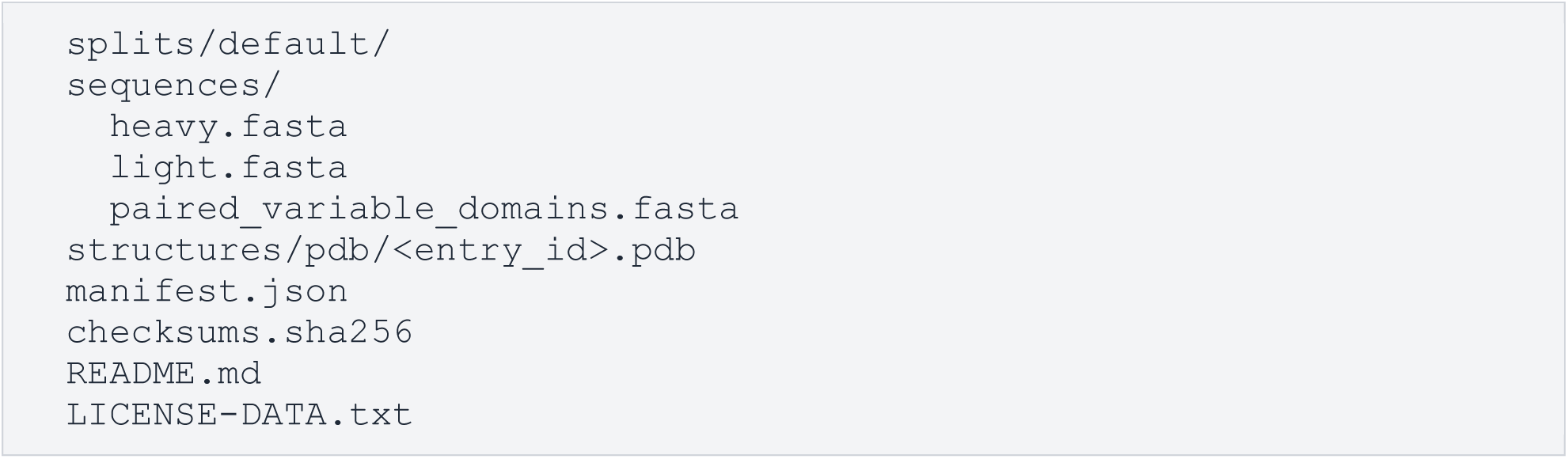

## Data Overview

The release contains 20,112 processed entries (Table 3): 16,511 experimental training entries, including 5,668 exact-CDR representatives; 3,274 synthetic patent-derived modelled complexes; and 327 held-out experimental entries. H3 length distributions are broadly similar between the non-redundant experimental training and test sets, while the patent-derived modelled collection contains a somewhat different distribution of loop lengths (Fig. 2A). The experimental collections contain diverse antigen types, whereas the patent-derived modelled component consists exclusively of protein antigens (Fig. 2B). Paired antibodies are more common than nanobodies in the experimental training and test sets, while the modelled collection contains comparable numbers of the two antibody types (Fig. 2C). Most experimental structures were determined by X-ray crystallography, with cryo-EM representing a smaller fraction, whereas all patent-derived structures were modelled with AlphaFold3 (Fig. 2D). Most exact-CDR redundancy groups contain only a single entry, with progressively fewer large clusters (Fig. 2E). Experimental training structures span several decades of PDB deposition, whereas the held-out test set is restricted to structures released after the 30 September 2021 cutoff (Fig. 2F). Test structures satisfy the ≤3.0 Å resolution criterion by construction, while the experimental training collection spans a broader range of resolutions (Fig. 2G). Missing CDR residues occur in about 13% of non-redundant experimental training structures, with their frequency varying by experimental method (Fig. 2H).

**Figure 2.**
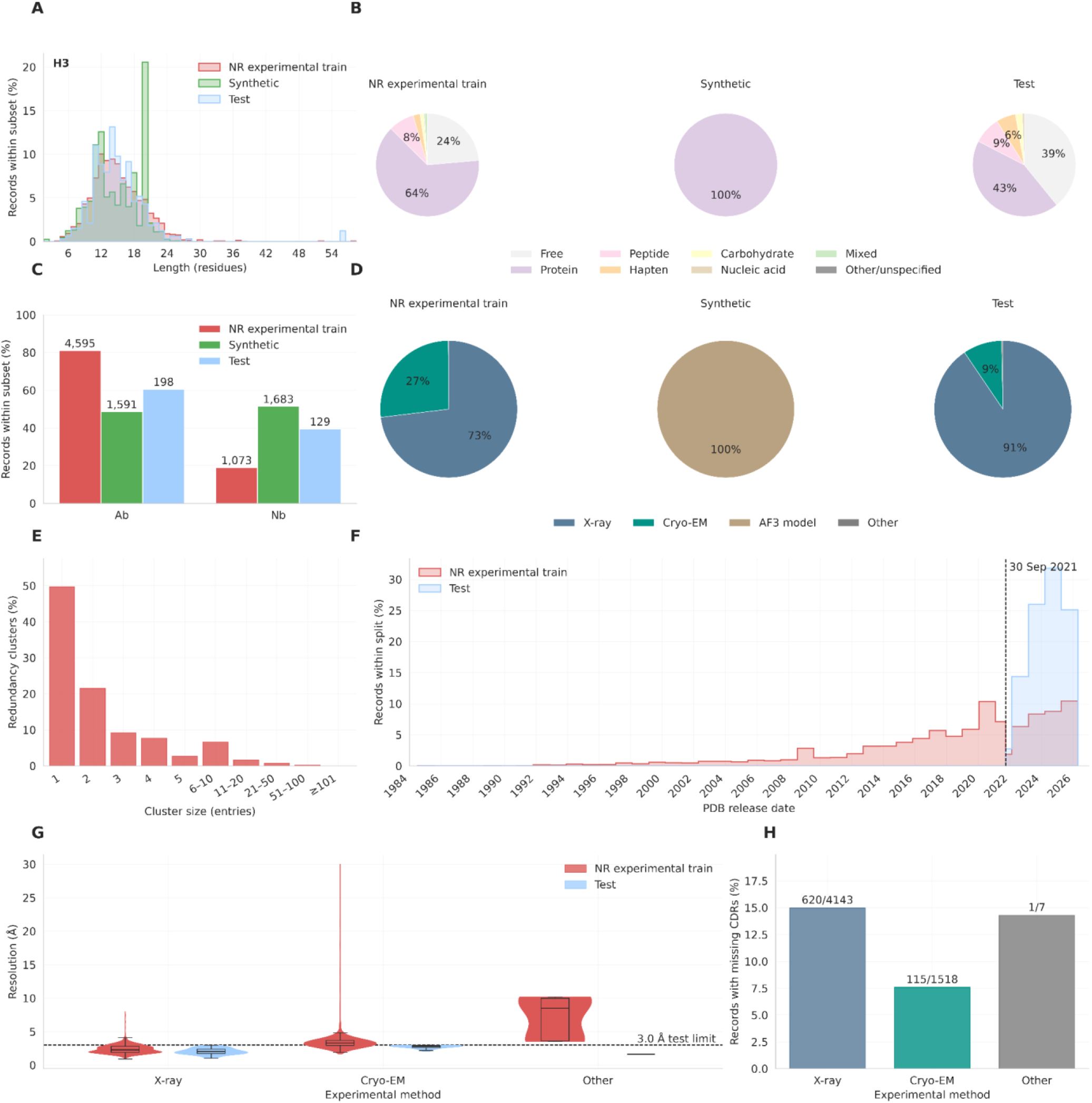
Dataset overview. **(A)** H3 length distributions. **(B)** Antigen-type composition. **(C)** Paired-antibody and nanobody proportions. **(D)** Experimental structure-determination or modelling method. **(E)** Exact-CDR redundancy-cluster sizes for experimental training data. **(F)** Experimental PDB release-date distributions; the vertical line marks 30 September 2021. **(G)** Reported resolution by experimental method for non-redundant training and test entries; the dashed line shows the 3.0 Å test set threshold. **(H)** Frequency of missing CDR residues among non-redundant experimental training representatives.

**Table 3.** Principal SABLE v0.2.0 subsets.

| Subset | Selection | Entries | Has Antigen |
| --- | --- | --- | --- |
| Experimental training - all | `split=train; synthetic=False` | 16,511 | 13,562 |
| Experimental training - non-redundant | `split=train; synthetic=False; non_redundant=True` | 5,668 | 4,331 |
| Patent-derived modelled | `split=train; synthetic=True` | 3,274 | 3,274 |
| Held-out test | `split=test` | 327 | 199 |
| Complete release | All rows | 20,112 | 17,035 |

## Technical Validation

### Confidence characteristics of the patent-derived models

All 3,274 patent-derived complexes passed the antigen-focused interface-confidence (ipTM) threshold by construction (Fig. 3A). Despite the strict antigen-focused ipTM cutoff (≥0.8), some retained complexes have relatively low antigen pTM values. Inspection indicated that these cases predominantly involved multi-domain proteins or proteins containing disordered regions where the antibody binds to a domain that was confidently modelled (Fig. S1). CDR pLDDT values were lower than framework pLDDT values, with H1 and H3 having the lowest confidence (Fig. 3B). Confidence varied with loop length: mean H3 pLDDT decreased with H3 length (Spearman’s ρ = -0.52, P = 2.1 × 10^-223; Fig. 3C), while ipTM had a weaker negative association (ρ = -0.22, P = 4.3 × 10^-37; Fig. 3D). These scores are technical indicators of predicted-model confidence^58^, not experimental evidence of binding, affinity, or biological activity. The released “synthetic” flag and confidence fields allow users to stratify analyses by loop length or confidence.

**Figure 3.**
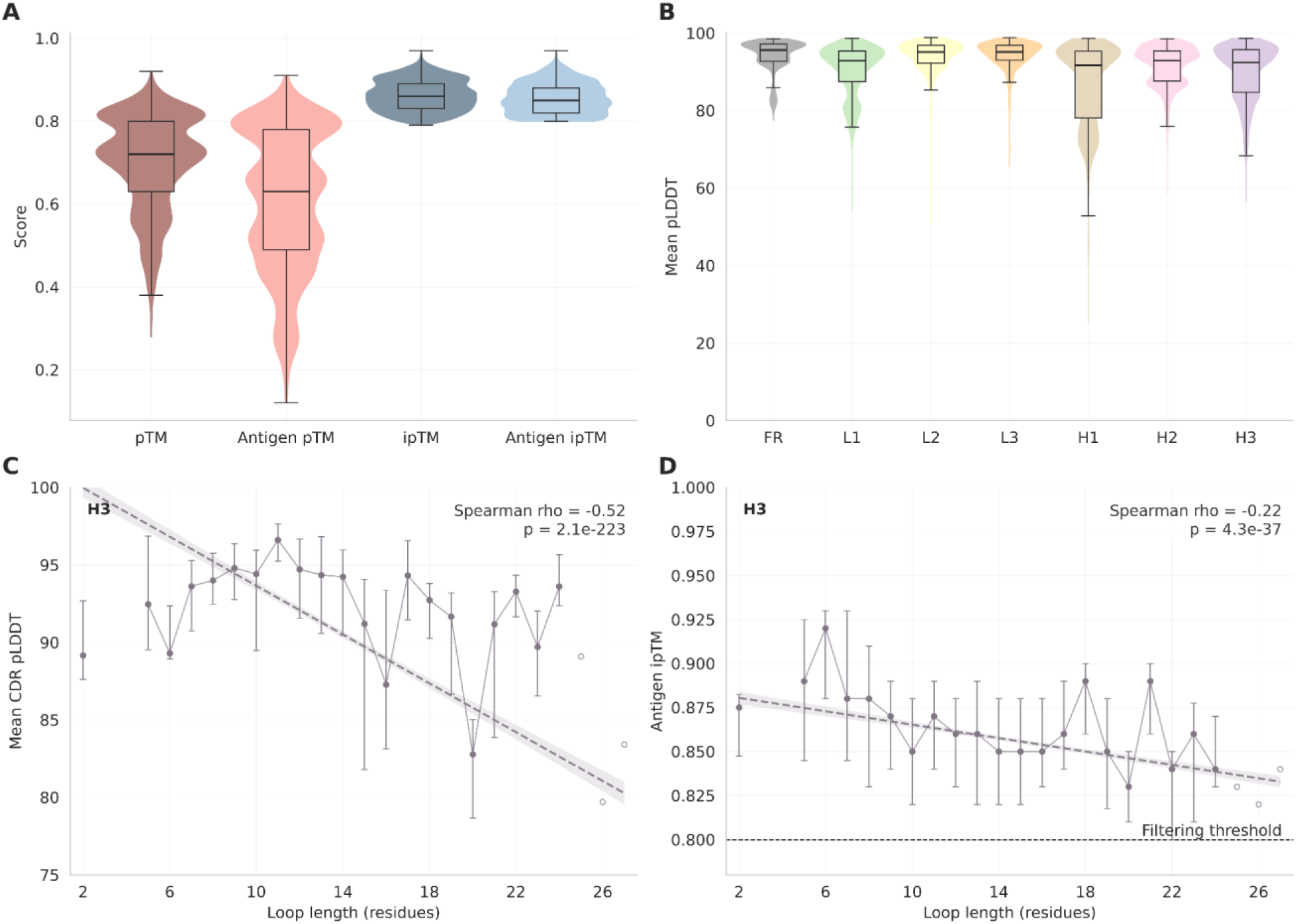
Confidence validation of the synthetic patent-derived modelled component. **(A)** Distributions of pTM, antigen-focused pTM, ipTM and antibody-antigen ipTM (“antigen_ptm” and “antigen_iptm”). **(B)** Mean pLDDT distributions. **(C)** Mean all-atom H3 pLDDT by loop length. Filled points summarise length groups with at least five entries as the median and interquartile range; open points represent rarer lengths. The dashed fit and band show the fitted trend and confidence interval. **(D)** Antigen-focused ipTM by H3 length, with the nominal retention threshold of 0.8. Spearman correlations and sample sizes are shown in the panels.

### Validation of train and test separation

The released nearest-neighbour fields make the split criteria directly auditable. We compared each test set entry with its experimental-training and patent-derived nearest neighbours. Every H3 distance is at least 3 (Fig. 4A) and every variable-domain identity is at most 90% (Fig. 4B), by construction. For the 170 test entries with protein or peptide antigens, the best-ranked BLAST hit remains below the unconditional 70% identity exclusion rule (Fig. 4C). For context, we also calculated leave-one-out nearest-neighbour baselines within the non-redundant experimental training set. We randomly sampled 327 training representatives for the antibody-based measures and, for each sampled entry, temporarily excluded that entry itself and searched the remaining non-redundant training collection for its minimum H3 distance and maximum variable-domain sequence identity. For the antigen baseline, 152 protein/peptide-antigen training entries were sampled and queried by BLAST against the non-redundant training antigen database, with self-hits excluded and the best remaining hit selected using the same bit-score, percent-identity, coverage and E-value ranking used for the released nearest-neighbour metadata. The mean of these leave-one-out nearest-neighbour values is shown as the grey baseline in Fig. 4A-C, providing a reference for the degree of similarity normally observed between distinct entries within the training collection and to validate the selected train-test split thresholds.

**Figure 4.**
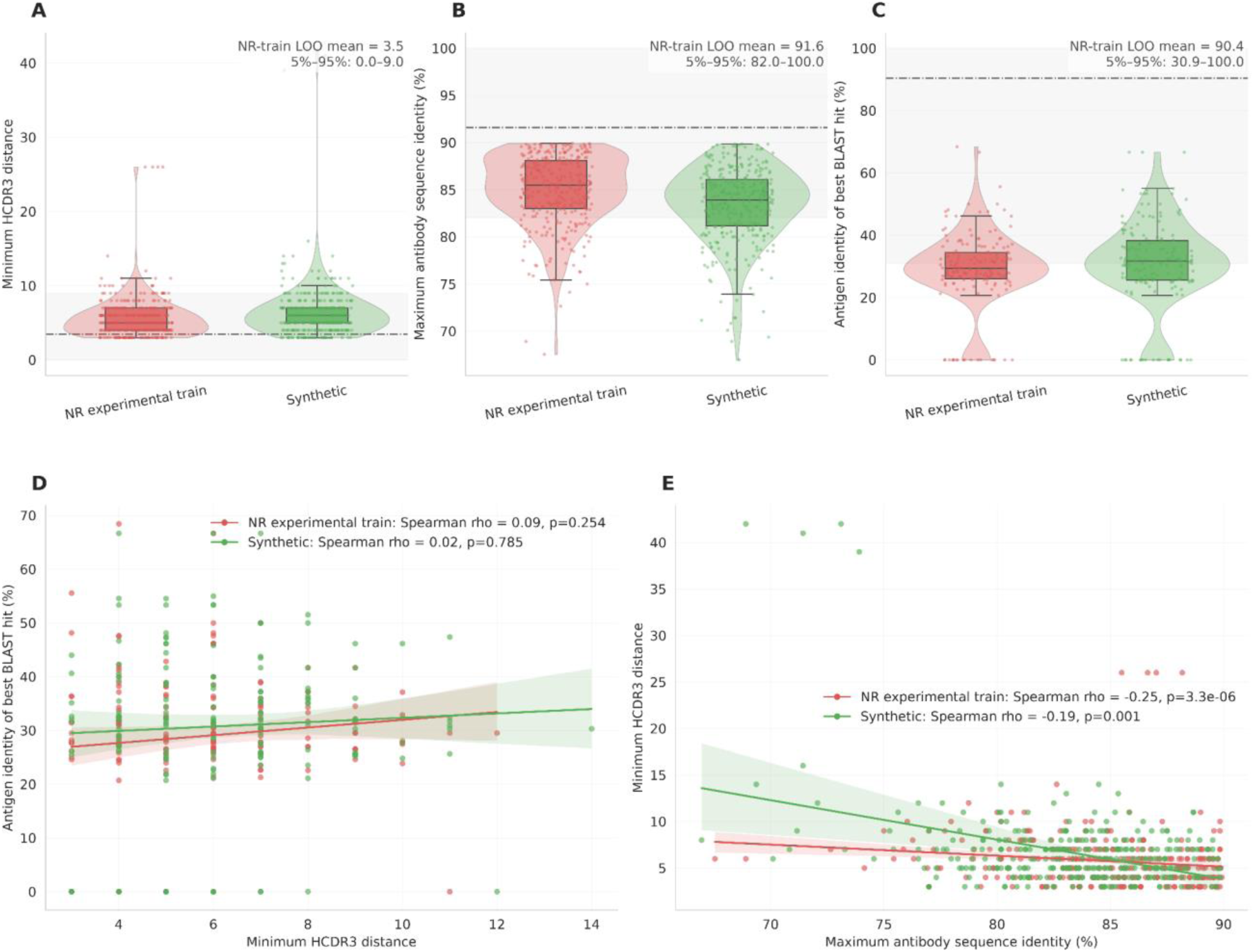
Nearest-neighbour validation of the held-out test set. **(A)** Minimum LCS-derived H3 distance from each test entry to experimental-training and patent-derived modelled references; the grey dash-dot line shows the sampled non-redundant-training leave-one-out baseline. **(B)** Maximum variable-domain sequence identity, with the leave-one-out baseline. **(C)** Percent identity of the best-ranked antigen BLAST hit for the 170 protein/peptide-antigen test entries; entries with no hit are represented as 0; the grey line shows the leave-one-out baseline. **(D)** H3 distance versus antigen BLAST-hit identity. **(E)** Maximum antibody identity versus H3 distance. Points are test entries; colours indicate the reference collection; lines and bands show trend estimates and confidence intervals.

H3 distance and antigen-hit identity showed weak correlations for both the experimental and modelled sets (experimental training: ρ = 0.09, P = 0.254; modelled: ρ = 0.02, P = 0.785; n = 170 each; Fig. 4D), indicating that the antibody and antigen controls capture different aspects of relatedness. H3 distance and maximum variable-domain identity showed modest inverse correlations (experimental training: ρ = -0.25, P = 3.3 × 10^-6; modelled: ρ = -0.19, P = 0.001; n = 327 each; Fig. 4E), supporting the use of both antibody-level criteria rather than substituting one for the other.

### Benchmark evaluation protocol

To demonstrate use of the predefined test set, we evaluated top-ranked complex predictions from AlphaFold3, Boltz-1 and OpenDDE^52,53,55^. These methods were selected because their reported training-data cutoffs are the same as the SABLE held-out period, thereby avoiding temporal overlap with the test set. For each method and test entry, the highest-ranked available structure was selected using the accompanying ranking or confidence score.

Region-level accuracy was measured from metadata-indexed Cα coordinates. For each heavy or light chain, predicted and native framework coordinates from FR1-FR4 were superposed with the Kabsch algorithm^59^, and CDR RMSD was calculated after this framework fit. Residues were paired by sequence alignment so that insertions or unresolved positions did not shift downstream coordinates. Antibody-antigen interfaces were evaluated with DockQ v2.1.3^27^ using the CAPRI protein-peptide protocol for peptide antigens^60^. Sequence-paired heavy and light chains were merged as the receptor, and all protein/peptide antigen chains were merged as the ligand before DockQ evaluation. For ambiguous or symmetry-related multichain antigens, sequence-compatible one-to-one antigen assignments were generated. Each mapping was evaluated and the mapping with the highest grouped DockQ score was retained. DockQ quality categories were incorrect (<0.23), acceptable (0.23 ≤ DockQ < 0.49), medium (0.49 ≤ DockQ < 0.80), and high (DockQ ≥ 0.80). Panels reporting top-1 success treat acceptable or higher quality models as successful. OpenDDE did not produce predictions for three test entries because of out-of-memory errors, including on an H200 GPU with 140 GB memory. The three failed OpenDDE runs were counted as failures in top-1 success-rate calculations but treated as missing for analyses of continuous DockQ distributions and correlations.

Prediction confidence was read from each method’s output files. Interface predicted TM score (ipTM) was compared with DockQ for protein/peptide antigen entries, and mean all-atom H3 pLDDT was compared with framework-aligned H3 RMSD. These evaluations were not used to choose records or construct the release; they provide a reproducible technical demonstration of how SABLE supports both region-level antibody assessment and interface-level complex assessment.

### Benchmark evaluation demonstration

We applied the SABLE evaluation API to the top-ranked AlphaFold3, OpenDDE and Boltz-1 predictions. Framework-aligned CDR RMSDs were computed for 327 AlphaFold3 and Boltz-1 predictions and 324 OpenDDE predictions. As expected, H3 had the broadest RMSD distribution (Fig. 5A). Mean H3 RMSD was higher for longer loops and for test entries farther from the nearest experimental-training H3 (Fig. 5B,C), demonstrating that these released metadata fields can be used for stratified evaluation.

**Figure 5.**
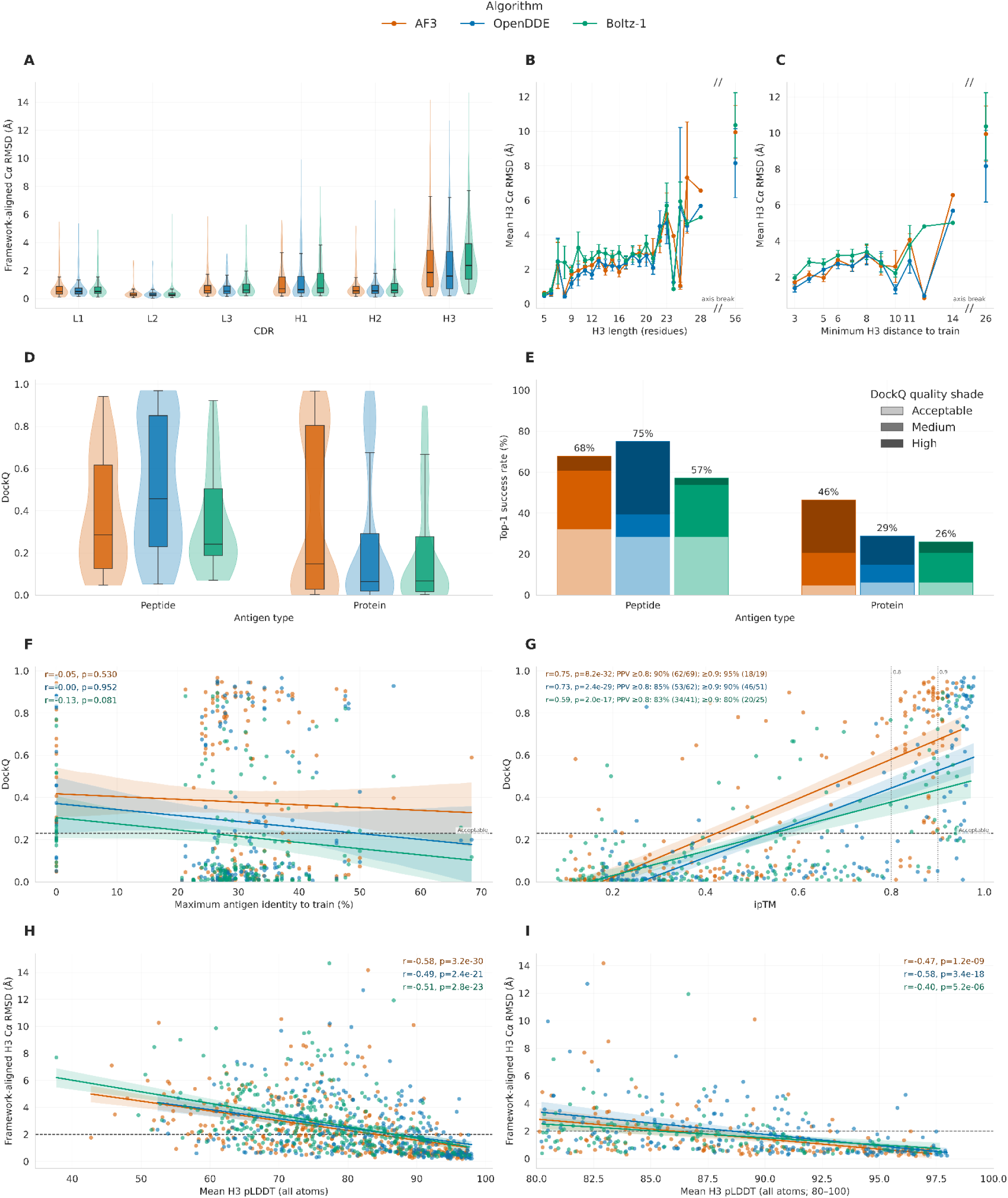
Benchmark evaluation of AlphaFold3, OpenDDE and Boltz-1. **(A)** Framework-aligned Cα RMSD for the six CDRs, calculated after superposition of the corresponding heavy- or light-chain framework regions. **(B)** Mean H3 RMSD by H3 length. **(C)** Mean H3 RMSD by minimum H3 distance to experimental training data. Points in B and C are group means with error bars. **(D)** DockQ distributions for peptide and protein antigens. **(E)** Top-1 acceptable or higher quality success rates, partitioned into acceptable, medium and high DockQ categories. **(F)** DockQ versus maximum antigen sequence identity to experimental training data; no BLAST hit values are represented as 0. **(G)** DockQ versus model-reported ipTM. Vertical lines mark ipTM 0.8 and 0.9, and panel text reports positive predictive value (PPV) for DockQ ≥ 0.23 at these thresholds. **(H)** Framework-aligned H3 RMSD versus mean all-atom H3 pLDDT. **(I)** The same relationship, restricted to the pLDDT range of 80-100. Dashed horizontal lines mark DockQ 0.23 in F and G and H3 RMSD 2 Å in H and I.

Approximately half of the test entries contained a protein (N=142) or a peptide (N=28) antigen. Acceptable-quality or higher top-1 rates for peptide antigens were 68%, 75% and 57%, for AlphaFold3, OpenDDE and Boltz-1, respectively, and 46%, 29% and 26% for protein antigens (Fig. 5D,E). DockQ showed little monotonic association with maximum antigen sequence identity to training (Fig. 5F), whereas model-reported ipTM was positively associated with DockQ for all three methods (Fig. 5G). At ipTM ≥ 0.8, the positive predictive value for DockQ ≥ 0.23 was 90% (62/69) for AlphaFold3, 85% (53/62) for OpenDDE and 83% (34/41) for Boltz-1; at ipTM ≥ 0.9, it was 95% (18/19), 90% (46/51) and 80% (20/25), respectively.

Mean all-atom H3 pLDDT was inversely correlated with H3 RMSD (AlphaFold3: ρ = -0.58; OpenDDE: ρ = -0.49; Boltz-1: ρ = -0.51; Fig. 5H). The inverse relationship remained within the high-confidence pLDDT range of 80-100 (ρ = -0.47, -0.58 and -0.40, respectively; Fig. 5I). These analyses provide a reproducible technical check of the evaluation workflow and confidence fields and are not intended as a ranking of the three methods.

## Usage Notes

### Training new deep-learning models

The intended benchmark pattern is to train or tune only on an explicitly declared training subset and reserve the predefined experimental test set for final evaluation. Users should report whether they used the full experimental training collection, the 5,668 exact-CDR representatives, the 3,274 patent-derived modelled entries, or a combination. Patent-derived structures may increase structural coverage but are confidence-filtered predictions rather than experimental observations; analyses should preserve the “synthetic”, “method” and model-confidence fields. Because this component contains AlphaFold3 outputs, downstream use is also subject to the applicable AlphaFold3 Output Terms of Use. Users should review the current terms before reuse, particularly for commercial use or training biomolecular structure-prediction models.

### Benchmarking new deep-learning models

Every reported SABLE benchmark result should identify the SABLE release version. The nearest-neighbour columns support task-specific audits, stricter relatedness thresholds and stratified reporting without changing the predefined split labels. Where relevant, benchmark results should be reported together with these relatedness measures.

Users who reproduce or extend the H3 distance or variable-domain identity calculations should use the SABLE API functions rather than substitute a generic distance or local-alignment implementation. Protein/peptide antigen neighbour fields follow the documented BLAST ranking, so “closest_*_blast_pident” should be interpreted together with bit score, E-value and coverage.

Predictions can be evaluated with the SABLE API to reduce implementation-dependent differences between studies. The “evaluate_antibody_prediction” workflow calculates the same sequence-mapped, framework-aligned CDR RMSDs used here and can additionally run the SABLE DockQ workflow for antibody-antigen complexes. Using these shared implementations keeps residue pairing, missing-residue handling, framework alignment, chain mapping and multimeric-antigen handling consistent with the benchmark definition.

### Example Python/PyTorch usage

A typical workflow can download the versioned release, select a benchmark subset and then access antibody-specific metadata, sequences, structures and coordinates directly:

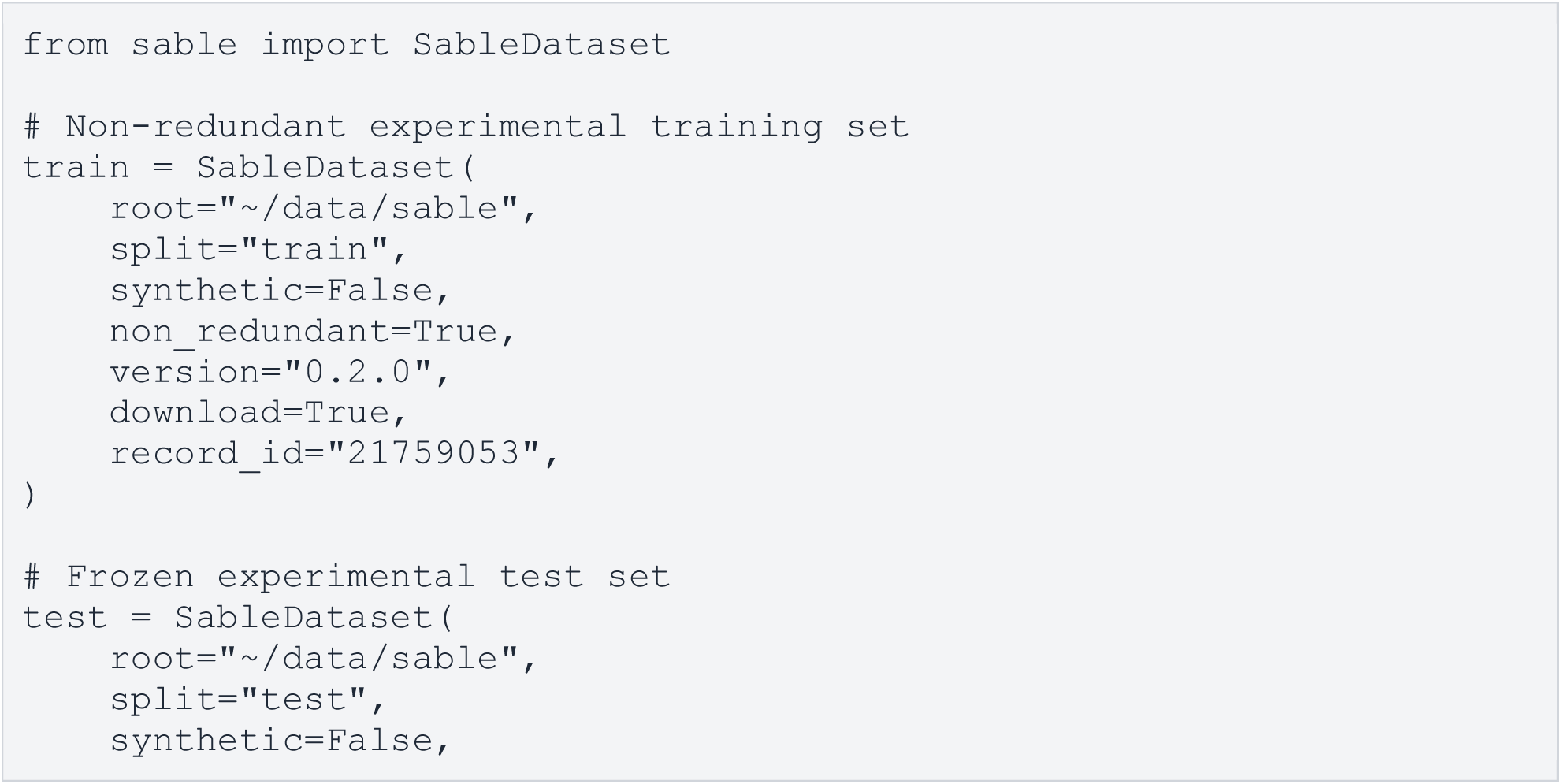
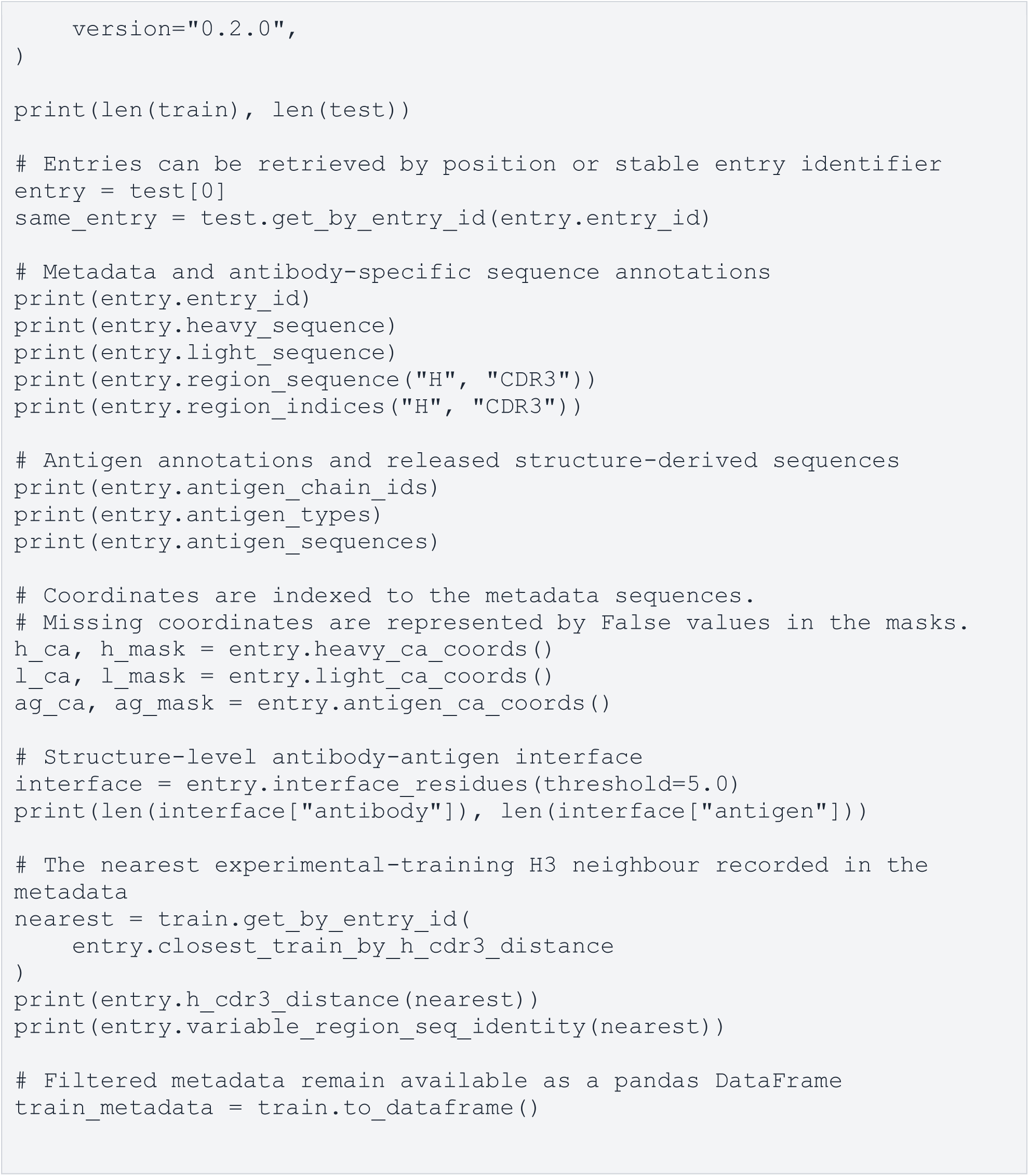

“SableDataset” can additionally filter entries by experimental or modelled origin, redundancy, antigen presence and protein/peptide-antigen availability, while “get_by_entry_id”, “iter_antibodies” and “to_dataframe” provide alternative access patterns for downstream analyses.

### Limitations of the dataset

SABLE reduces documented antibody- and antigen-sequence overlap but cannot eliminate every form of biological relatedness. Antigens with low sequence identity can retain remote structural homology, functionally related antibodies can have different H3 sequences, and hapten, carbohydrate and nucleic-acid antigens are not screened with the same BLASTP procedure as protein/peptide antigens. Each processed entry is defined by the source coordinate model and SAbDab chain annotation rather than an independently generated biological assembly. Users whose tasks depend on oligomeric state, alternative assemblies, remote structural similarity or binding measurements should apply additional checks using the released provenance and structure files.

## Data Availability

SABLE v0.2.0 metadata and dataset components not subjected to third-party restrictions are available from Zenodo under DOI 10.5281/zenodo.21759053^57^ and released under the Creative Commons Attribution 4.0 International licence. Patent-derived AlphaFold3-generated structure outputs are subject to the applicable AlphaFold3 Output Terms of Use at: https://github.com/google-deepmind/alphafold3/tree/main. The record contains CSV and Parquet metadata, the JSON schema and column dictionary, predefined split and redundancy files, sequence FASTA archives, sharded processed-PDB archives, a manifest, checksums, a validation report, README and data licence. An interactive browser is available at https://bio3d.cs.huji.ac.il/webserver/sable/.

## Code Availability

The SABLE Python package, PyTorch-compatible dataset loaders, validation tests and benchmark metrics are available under the MIT licence at https://github.com/dina-lab3D/SABLE. Version 0.2.0 provides commands for downloading and verifying the Zenodo record, safe extraction, metadata filtering, structure and coordinate access, antigen-sequence handling, LCS-derived similarity, framework-aligned CDR RMSD and DockQ v2.1.3 evaluation. Downstream users should record the package version and Git commit used.

## Author Contributions

T.C. conceived and led dataset construction, implemented the processing, validation and software workflows, performed the analyses and drafted the manuscript. H.B. contributed to data curation, documentation and web resource development. M.O.-F. contributed to the design and curation of the patent-derived modelled component and revised the manuscript. D.S. supervised the project, contributed to methodology and revised the manuscript. All authors reviewed and approved the manuscript.

