## Supplementary for "A structural antibody benchmark for leakage-aware deep-learning evaluation"

**Table S1 | Test set manual inspection**

| Entry ID | Decision | Notes |
|---|---|---|
| 7y8j_0 | Remove | COVID-19 peptide |
| 8cge_0 | Remove | Fused antibody |
| 8dqu_0 | Change antigen stoichiometry | Dimer antigen |
| 8dtu_0 | Change chain IDs | Reassigned antigen chain from B to A |
| 8ee2_1 | Change antigen stoichiometry | Dimer antigen |
| 8eq6_0 | Remove | Contains an antigen that is not mentioned in SAbDab metadata. |
| 8fif_0 | Remove | Microcystin-LR peptide |
| 8j7t_0 | Change antigen stoichiometry | Dimer antigen |
| 8p88_0 | Remove | A light chain with a nanobody complex |
| 8r3w_0 | Remove | Diabody |
| 8rpe_0 | Remove | A light chain with nanobody complex |
| 8s72_0 | Change antigen stoichiometry | Dimer antigen |
| 8v91_0 | Change antigen stoichiometry | Tetramer antigen |
| 8v9d_0 | Change antigen stoichiometry | Tetramer antigen |
| 8vyl_1 | Change antigen stoichiometry | Tetramer antigen |
| 9b2w_0 | Change antigen stoichiometry | Dimer antigen |
| 9egn_0 | Change antigen stoichiometry | Trimer antigen |
| 9g2g_1 | Change antigen stoichiometry | Dimer antigen |
| 9gvn_0 | Change antigen stoichiometry | Dimer antigen |
| 9luy_0 | Remove | Fused peptide antibody |
| 9pym_0 | Remove | Megabody |
| 9uln_0 | Remove | Same antibody as 8v9d_0 without an antigen. |
| 8v9d_1 | Remove | Same antibody as 8v9d_0 with monomer antigen instead of a dimer. |
| 9x1h_0 | Change antigen stoichiometry | Dimer antigen |

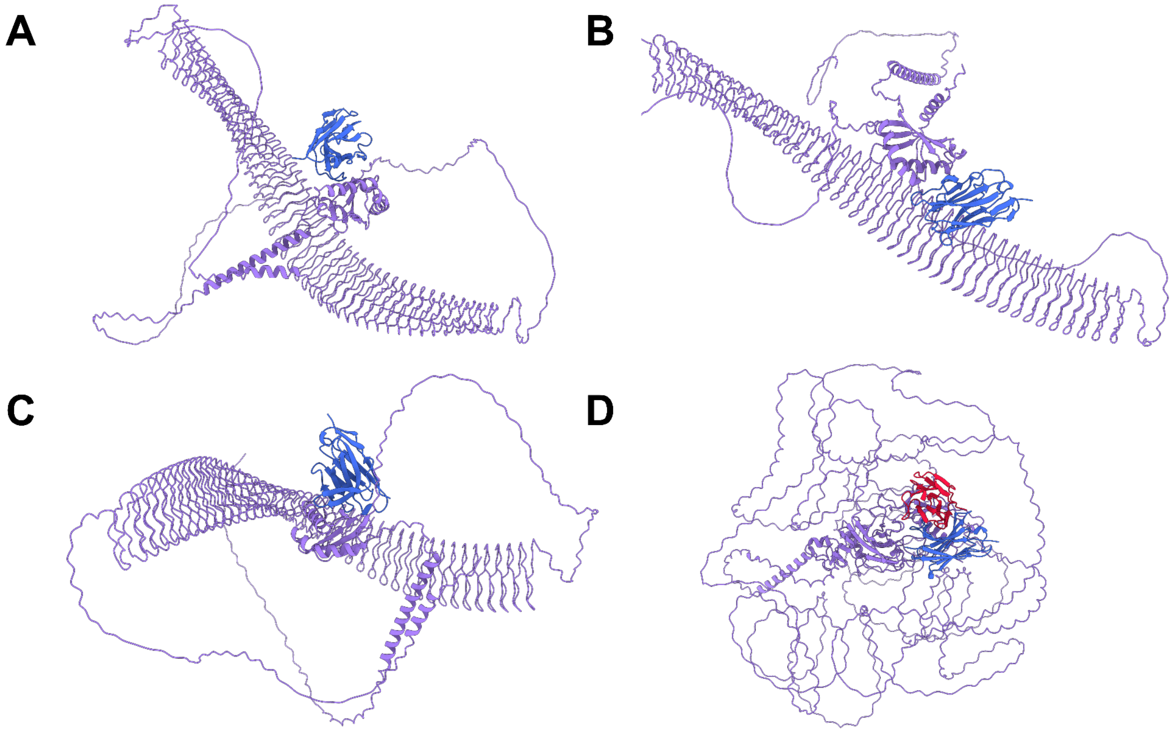

**Figure S1 | Patent-derived entries with the lowest pTM values. Antigen chains are shown in purple, heavy chains in blue and light chains in red. (A)** Entry p15941_214812 (pTM=0.28). **(B)** Entry p15941_169017 (pTM=0.28). **(C)**. Entry p15941_111391 (pTM=0.28) **(D)** Entry p02452_139974_2 (pTM=0.3).
